# 3D chromatin interactions involving *Drosophila* insulators are infrequent but preferential and arise before TADs and transcription

**DOI:** 10.1101/2022.12.12.520036

**Authors:** Olivier Messina, Flavien Raynal, Julian Gurgo, Jean-Bernard Fiche, Vera Pancaldi, Marcelo Nollmann

## Abstract

In mammals, insulators contribute to the regulation of loop extrusion to organize chromatin into topologically associating domains. In *Drosophila* the role of insulators in 3D genome organization is, however, under current debate. Here, we addressed this question by combining bioinformatics analysis and multiplexed chromatin imaging. We describe a class of *Drosophila* insulators enriched at regions forming preferential chromatin interactions genome-wide. Notably, most of these 3D interactions do not involve TAD borders. Multiplexed imaging shows that these interactions occur infrequently, and only rarely involve multiple genomic regions coalescing together in space in single cells. Finally, we show that non-border preferential 3D interactions enriched in this class of insulators arise before TADs and transcription during *Drosophila* development. Our results are inconsistent with insulators forming stable hubs in single cells, and instead suggest that they fine-tune existing 3D chromatin interactions, providing an additional regulatory layer for transcriptional regulation.

## Introduction

Eukaryotic chromosomes are organized in a multi-layered structure comprising chromosome territories, compartments, topologically-associating domains (TADs) and nano-domains ^1,2^. Notably, this multi-scale organization of the genome is conserved from *Drosophila* to mammals ^3–6^. However, the mechanisms responsible for the acquisition and maintenance of these structures seem to differ between species.

In vertebrates, TADs are often separated from each other by convergent CCCTC-binding factor (CTCF) sites localized at TAD boundaries. TAD borders bound by CTCF/cohesin form “focal chromatin loops” in contact matrices ^7^. These specific looping interactions may facilitate the communication between genes and their cis-regulatory elements (CREs, e.g. enhancers and promoters) most often localized within TADs ^8,9^. In mammals, the formation of TADs is thought to involve loop extrusion, a mechanism by which Structure Maintenance of Chromosome (SMC) proteins (e.g. cohesin) bind chromatin and reel it in until they encounter properly-oriented CTCF sites ^10^. In contrast, the *Drosophila* homologue of CTCF (dCTCF) binds preferentially within TADs and is only mildly enriched at TAD borders ^4,6^ with no preferential convergent orientation as in mammals ^11^. These results suggest that other mechanisms may instead be responsible for the establishment of TADs in *Drosophila*.

While CTCF is the main chromatin insulator in mammals, *Drosophila* contains tens of non-evolutionary conserved insulator binding proteins (hereafter IBPs) ^12^. Since before the genomic era, *Drosophila* insulators were shown to be involved in the regulation of long-range chromatin interactions: either by blocking enhancer-promoter interactions or by establishing barriers between chromatin states ^13–15^. Early genome-wide studies showed that insulators preferentially bind to genomic regions containing housekeeping genes and highly transcribed regions ^16^. In addition, IBPs frequently bind to TADs borders ^4,6,17–19^ that can often interact in 3D ^20^. Taken together, these data suggest that insulators may be involved in the organization of *Drosophila* TADs, yet do not seem to rely on loop extrusion.

Recent studies suggest different modes of action. On one hand, insulators may promote TAD border interactions by forming long-range contacts between insulator factors ^21–23^. On the other hand, insulators may not form CTCF-like focal chromatin loops, but rather restrict interactions between domains ^24^.

Here, we investigated the role of insulators in the 3D organization of the *Drosophila* genome by combining advanced bioinformatics analysis and Hi-M, an imaging-based method we recently developed to detect the 3D positions of multiple genomic loci in single cells ^25^. First, we show that genomic regions occupied by insulators display a preference to co-localize in space. This preferred co-localization occurs inside TADs, can span TAD borders, and sometimes implicates multiple chromatin loci. Second, we show that chromatin regions displaying the most prominent 3D interactions are preferentially bound by insulators. We detect TAD border preferential interactions, but these represent the minority of the interactions detected. Interestingly, non-border interactions quantitatively increased with the occupancy of IBPs. Finally, by mapping these interactions during development, we found that non-border regions harboring insulators interact in 3D before the emergence of TADs.

## Results

### Genomic regions displaying preferential interactions are predominantly bound by chromatin insulators

To shed light onto the roles of *Drosophila* insulators in 3D genome organization during early embryogenesis, we deployed Chromatin Assortativity analysis (ChAs) ^26,27^. Assortativity measures the preference for the nodes of a network to interact with other nodes that have the same characteristics. In ChAs analysis, a chromatin interaction network is built from a single genome-wide contact map. This network represents the genomic loci (nodes) displaying high frequency interactions (i.e. loops) amongst each other (**Fig. 1a**, see *chromatin assortativity* in Methods). Chromatin assortativity for a given factor is calculated by estimating whether nodes bound by this factor interact with other nodes with the same factor more than expected at random. Thus, a factor with positive assortativity is enriched in chromatin loci that preferentially interact.

**Figure 1.**
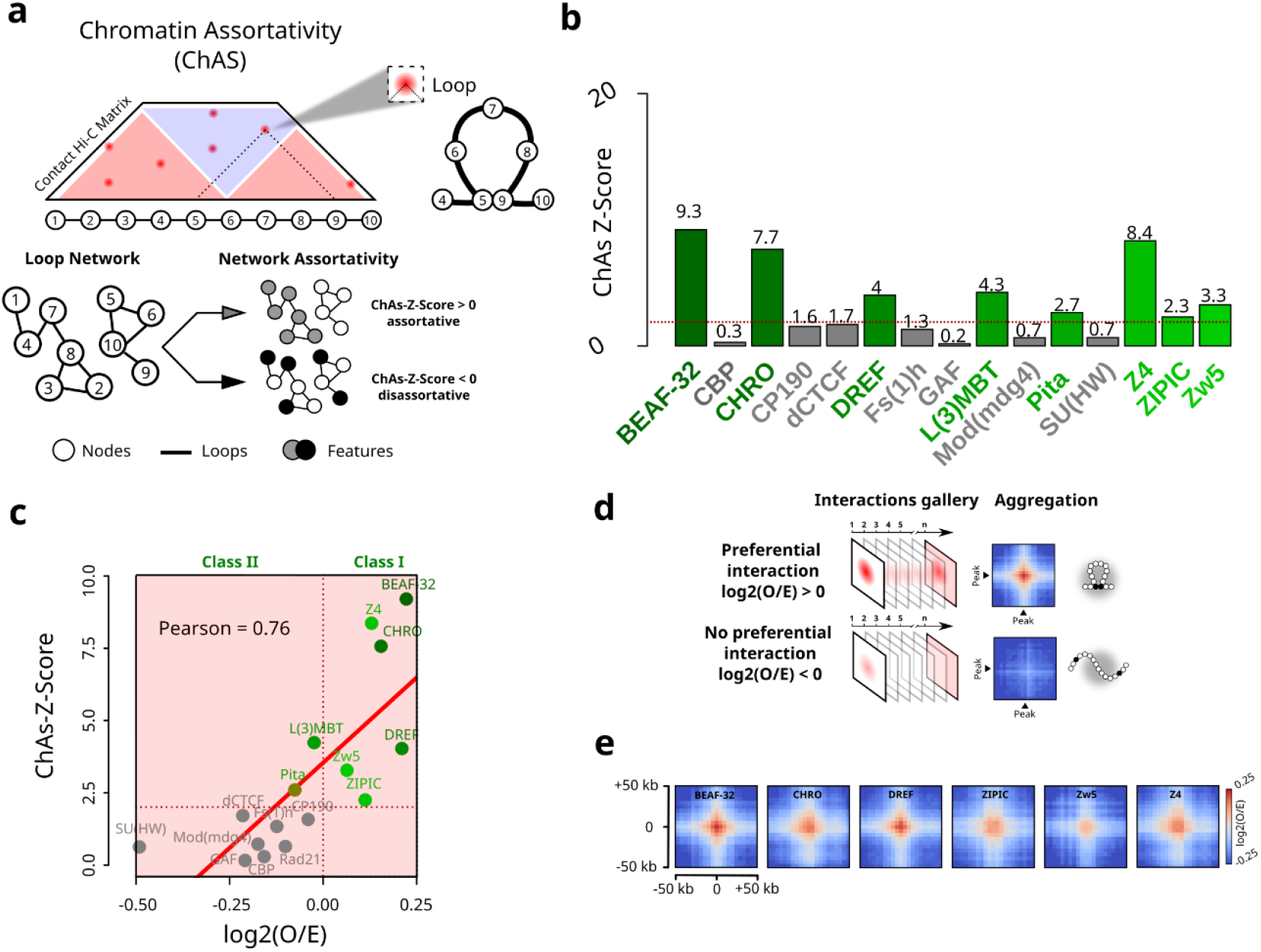
Genomic regions displaying preferential interactions are predominantly bound by chromatin insulators. a. Cartoon illustrating Chromatin Assortativity (ChAs) of chromatin binding factors in a network of chromatin contacts. 5-kb genomic bins are represented by nodes in the chromatin network. Nodes are connected to each other if they are making loops in Hi-C data (called by Chromosight [Matthey-Doret et al, 2020]). Nodes are color coded by the presence or absence of a given chromatin binding factor (features). The assortativity is then calculated for each different factor (see Methods). b. Barplot illustrating ChAs-Z-Scores for 15 IBPs in the nc14 chromatin network classified by alphabetical order. The horizontal red dash line represents the chAs-Z-Score=2 threshold considered in this study. c. Pearson’s correlation between ChAs-Z-Scores (y) and log2(O/E) (x) from APA for the 15 IBPs tested. Class I and II are delineated by a vertical red dash line centered in log2(O/E) = 0. d. Cartoon illustrating aggregate Hi-C map procedure around pairs of specific chromatin binding factors (peak). The first line illustrates the log2(O/E) aggregate map expected for a given factor involved in preferential contact formation and the second one a factor with no implication in the formation of preferential contacts (see Methods). e. Aggregate Hi-C plots of class I IBPs regions in nc14 embryos ^20^. Maps show the log2(O/E) in a 50 kb window around the crossing point of two class I IBPs regions : BEAF-32, DREF, CHRO, Z4, ZIPIC, Zw5.

We applied ChAs analysis to study chromatin organization of *Drosophila* embryos at nuclear cycle 14 (nc14) ^20^, a developmental stage coinciding with the zygotic genome activation (ZGA) and with the emergence of TADs ^20^. For this, we obtained chromatin interaction networks by mapping chromatin regions forming loops using Chromosight ^28^ on Hi-C data. Next, we annotated these chromatin networks with the binding patterns of publicly available ChIP-seq datasets (features, **Fig. 1a**) and calculated chromatin assortativities for a wide panel of chromatin binding factors, including insulator and insulator-associated proteins (BEAF-32, CBP, CHRO, CP190, dCTCF, DREF, FS(1)h, GAF, L(3)MBT, Pita, Mod(mdg4), Su(HW), Z4, ZIPIC and Zw5), pioneering factors (Zelda), RNA polymerase II (RNAPII CTD phospho-Ser5 : S5P), Polycomb group proteins (Pc, Ph) and the cohesin subunit (Rad21).

Chromatin assortativity Z-scores (hereafter ChAs Z-scores) are calculated to estimate if ChAs for a feature is higher than expected for regions separated by similar genomic distances, indicating the importance of 3D interactions for establishing preferential contacts. Regions enriched in Zelda, Polycomb group proteins (Pc and Ph), and RNAPII CTD phospho-Ser5 (S5P) displayed positive ChAs Z-scores (**Fig. S1a**), consistent with previous findings ^29–31^. In contrast, ChAs Z-scores were highly variable between IBPs (**Fig. 1b**), indicating that different insulators may contribute unequally to the formation of preferential contacts.

A sub-group of IBPs displayed high assortativities (ChAs Z-score > 2), including the insulator and insulator-associated proteins: BEAF-32, CHRO, DREF, L(3)MBT, Pita, Z4, ZIPIC and Zw5 (**Fig. 1b**). Notably, cohesin (Rad21), dCTCF, and a second sub-group of IBPs including CBP, CP190, Fs(1)h, GAF, Mod(mdg4) and SU(HW) displayed low assortativity and low Z-scores (ChAs Z-score < 2, **Figs. 1b, S1a**). The low assortativity of GAF may appear as surprising, as in the literature it was found that GAF is often present at focal loops (involving a few hundred loci) directly visible in Hi-C maps ^29,31^. Low assortativity scores can arise when all nodes of the network are either bound or not bound by a given factor, or when presence of this factor is not associated with presence of loops. As GAF is not present in all nodes, we conclude that most detected loops are not bound by GAF, even when a small subset of GAF sites may be associated with focal loops. Overall, these results indicate that *Drosophila* insulators tend to be associated with 3D interaction networks to different degrees, but do not shed light on whether and how binding of insulators may modulate interaction frequencies.

### Insulator binding increases the strength of preferential long-range chromatin contacts

To address these questions, we complemented ChAs with aggregation peak analysis (APA)^7^. This method relies on the calculation of pairwise, intra-arm autosomal contact frequencies between genomic regions bound by a given factor (i.e. peak) (**Fig. S1b**). The statistical relevance of these contacts is estimated by calculating the average of the log2(Observed/Expected) distribution of the Hi-C signal at all peak regions (*see aggregate peak analysis* in Methods). Thus, the log2(O/E) ratio is positive when contacts occur at frequencies higher than expected and is negative when contact frequencies are lower than expected for regions separated by the same genomic distance.

Notably, the positive correlation between ChAs and log2(O/E) (**Fig. 1c**) indicates that the chromatin factors that are bound to regions forming preferential interactions (positive ChAs Z-scores) most often increase the strength of the interaction (positive log2(O/E) scores). Remarkably, most of the insulator factors displaying positive ChAs Z-Scores also exhibited positive log2(O/E) (BEAF-32, CHRO, DREF, Z4, ZIPIC and Zw5) (hereafter referred to as class I insulators) (**Figs. 1b, S1c**). Thus, class I insulators occupy genomic regions displaying the highest preferential interactions, and conversely, the genomic regions they occupy tend to preferentially interact in 3D in nc14 embryos.

Next, we investigated the specificity of preferential chromatin interactions by using Hi-C aggregate plot analysis ^20^ (**Fig. 1d**, and *Hi-C aggregate plot analysis* in Methods). Class I IBPs displayed a well-defined center spot, indicating that presence of IBPs at both loop anchors reinforces preferential 3D interactions (**Fig. 1e**). Similar results were observed for Zelda and RNAPII (**Fig. S1e**). In contrast, factors with low assortativity and/or negative log2(O/E) did not exhibit centered spots (**Fig. S1d**), likely due to positive and negative log2(O/E) values for different regions averaging out. These findings suggest that class II IBPs are less systematically involved in the establishment of preferential contacts in nc14 embryos.

### Most insulator-bound preferential long-range contacts involve non-border chromatin regions

TAD borders in *Drosophila* are mostly occupied by insulators ^4,6^, with only 4% of borders lacking insulator sequences ^19^. In *Drosophila*, ensemble analysis showed that adjacent TAD borders tend to preferentially interact ^20^, however contact maps do not display focal chromatin loops (e.g. corner peaks) as those observed in mammals ^7,32,33^. To determine whether our unbiased network analysis was able to recover preferential interactions between TAD borders, we calculated whether TAD borders were assortative in the network of chromatin interactions generated by Chromosight for nc14 embryos. This analysis shows that TAD borders appear highly connected to each other in the interaction network (**Figs. 2a-b**). This connectivity results in considerably higher ChAs values for TADs borders (**Fig. 2b**, blue dashed line) as compared to randomized networks (**Fig. 2b**, black distribution). As expected, and consistent with previous analysis ^20^, TAD borders exhibited a well-centered interaction spot in Hi-C aggregate plots (**Fig. S2a**).

**Figure 2.**
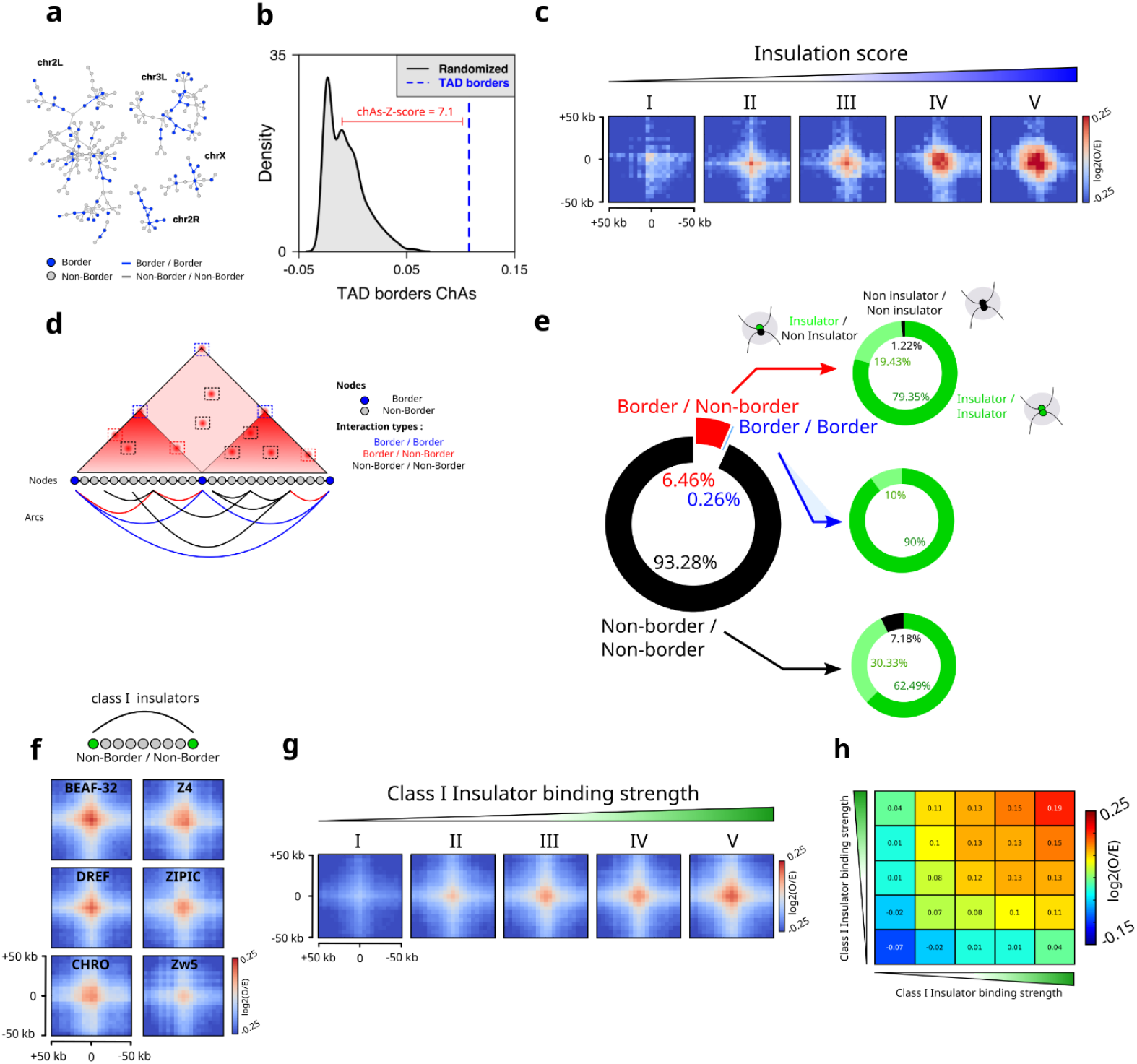
Border-Border interactions and non-border loops are favored by increase of insulation-score and IBPs binding. a. Chromosight chromatin network from Hi-C data at nc14 embryos ^20^. Each node of the network is a chromatin fragment, blue nodes represent nodes in which a TAD border is found, and edges represent significant 3D interactions. b. ChAs score for TAD borders (blue) versus distribution of ChAs scores for randomized networks (black). The ChAs-Z-Score is calculated between the distribution of the randomized network and the assortativity of TAD borders (see Methods). c. Aggregation Hi-C plots for TADs borders in nc14 stratified in five egal size categories groups with an increasing level of insulation score. d. Cartoon illustrating the different types of interaction observed in Hi-C dataset. Genomic bins are represented by color coded nodes. Arcs represent loops between pairs of genomic bins. Border / Border interactions are labeled in blue, Border / Non-Border in red and Non-Border / Non-Border in black. e. Pie plot representing the loops distribution called by chromosight into the different types of interaction (left panel). Pie plots illustrating the quantification of class I IBPs bound on each side of the anchored loop (right panel). f. Aggregation Hi-C plots for non-border regions bound by class I insulators in nc14 embryos. g. Aggregation Hi-C plots for class I IBPs in nc14 stratified in five egal size categories groups with an increasing level IBPs binding strength. h. Log2(O/E) between the different categories of class I insulator-bound in nc14 embryos.

The strength of a TAD border, as assessed by its insulation score (IS), is positively correlated to the binding level of insulator proteins ^19,34^. Given this correlation, we tested if the presence of IBPs at TAD borders is also associated with their interaction preference by stratifying TAD borders into five equally-sized categories according to their IS and by computing Hi-C aggregate plots for each category. Notably, the level of preferential interactions between TAD borders increased with insulation strength (**Fig. 2c**), providing indirect evidence for a role of IBPs in contributing to TAD border interactions.

Next, we wondered whether preferential chromatin contacts may be detected in locations other than TAD boundaries and what their determinants may be. To this aim, we divided the interactions in our chromatin network into three categories: border/border (blue), border/non-border (red), and non-border/non-border (black) (**Fig. 2d**). Then, we quantified the occurrence of each type of interaction by quantifying the presence of a border on each loop anchor. Notably, preferential interactions involved a border in one or both of their anchors in a minority of cases (<1% for border/border and ~6% for border/non-border), with the overwhelming majority of preferential interactions involving non-borders (>93%) (**Fig. 2e**).

To better understand the role of insulators in each of these interaction categories (i.e. border/border, border/non-border and non-border/non-border), we calculated for each category the proportion of interactions displaying binding of class I IBPs in two, one or none of the loop anchors. The vast majority of the anchors for all categories were bound by at least one class I IBPs (>92%, **Fig. 2e**, right pie charts). Anchors in border-border loops are most often bound by two class I IBPs (~90%), and in a smaller proportion by a single class I IBP. This trend was similar for the other categories, further supporting a role of class I IBPs in the mediation of chromatin loops that in most cases do not involve TAD borders.

To further support this conclusion, we performed aggregation Hi-C analysis on non-border/non-border loops. Notably, this analysis displays a strong aggregation peak, consistent with class I IBPs binding to both anchors of non-border interacting loops (**Fig. 2f**). Consistently, interactions mediated by class I IBPs at non-border regions increased with insulator binding strength (**Figs. 2g, S2b**) of both loop anchors (**Figs. 2h, S2c**). All in all, these analyses suggest that class I insulators participate in mediating preferential interactions between border and non-border chromatin regions. These analyses, however, do not inform us on how frequently these preferential interactions occur in single cells, or whether they involve two or multiple anchors.

### Insulator-bound chromatin regions only infrequently co-localize in 3D

Sequencing-based 3C methods only provide relative interaction frequencies, thus we turned to DNA-FISH, a technique that can quantify absolute physical proximity frequencies ^35,36^. As conventional DNA-FISH can only measure proximity between a limited number of genomic targets, we used Hi-M, a multiplexed imaging method that enables the detection of tens of genomic loci at once ^25,30,37^. Specifically, we imaged the 3D chromatin organization of the *dpp* locus (*chr2L: 2343645-2758688* dm6) in intact nc14 *Drosophila* embryos at ~12 kb resolution (**Figs. 3a, S3a**). The *dpp* locus contains three TADs and several regions displaying high levels of class I insulator binding, named barcode I1 to I10 (**Fig. 3b**). To cover this locus, we designed 34 equally-spaced barcodes that label insulator-bound and insulator-free genomic regions (**Fig. 3c**). Nuclei and barcodes were registered, segmented and localized as in previous studies ^25,30^ (**Fig. S3a**, see Methods), with similar barcode detection efficiencies (**Figs. S3c-d**). Ensemble pairwise distance maps were built by kernel density estimation of the full pairwise distance (PWD) distributions (**Fig. S3b**). Proximity maps were constructed by calculating the frequency of co-localization for each pair of barcodes from chromatin traces (**Fig. S3e**) using a pre-established distance threshold that maximizes the correlation between Hi-M and Hi-C datasets (d = 250 nm, **Figs. S3f-g**), and that was previously used for similar studies ^30^.

**Figure 3.**
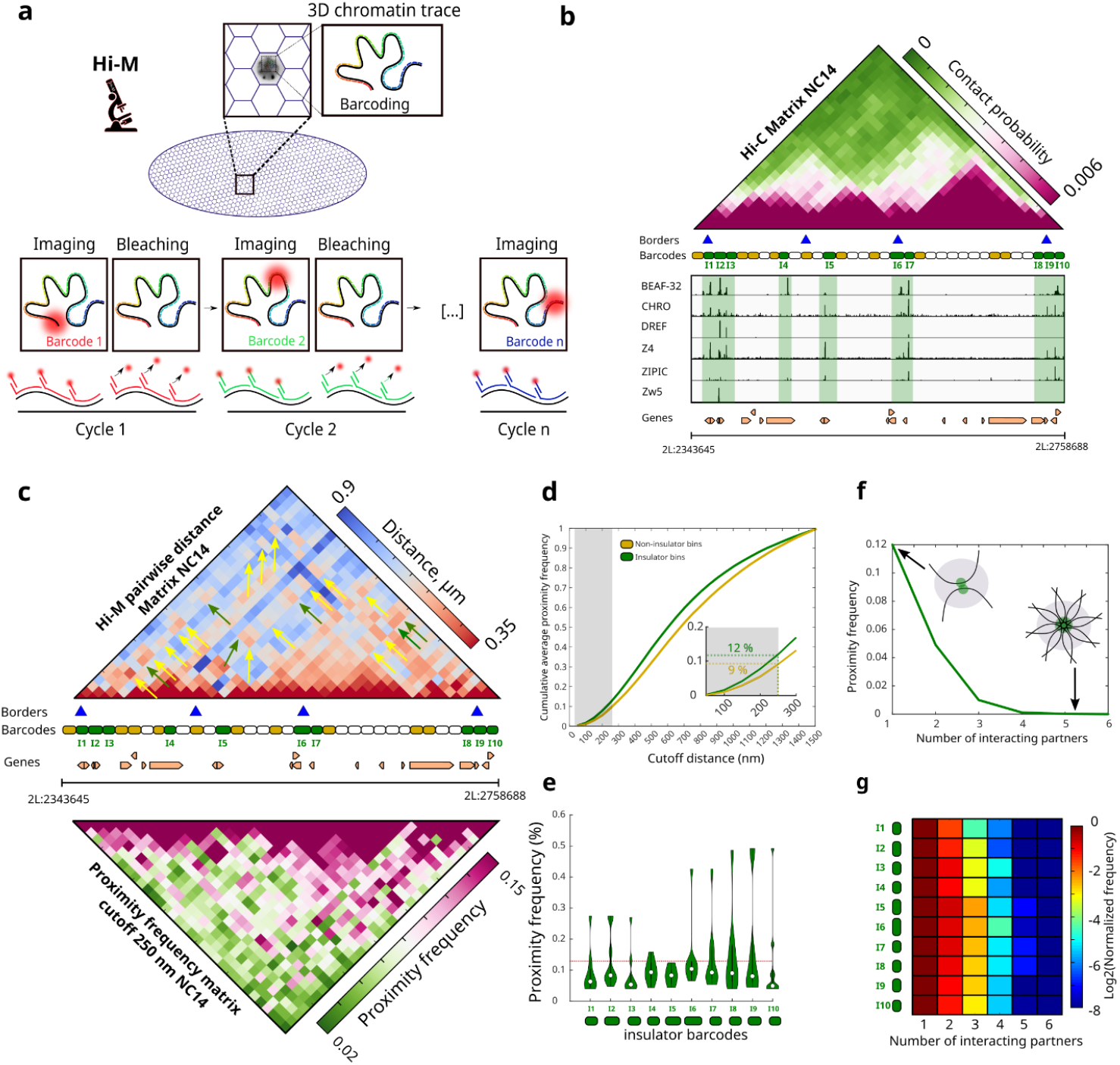
Hi-M reveals visible interactions between insulator-bound chromatin regions. a. Cartoon illustrating the imaging based strategy used to study chromosome conformation in intact drosophila embryos (Hi-M). b. Nc14 Hi-C matrix along the dpp locus (2L:2343645-2758688) in Drosophila melanogaster (dm6). Purple and green represent high and low contact probability respectively. Identified TADs borders from nc14 embryos (Hug et al. 2017) are represented by blue triangles. Barcodes used for Hi-M sequential imaging strategy are represented by color-coded rectangles. Barcodes bound by IBPs are filled in green and control regions in yellow. ChIP-seq profiles for class I IBPs (BEAF-32, CHRO, DREF, Z4, ZIPIC, Zw5) are displayed at the bottom together with genomic coordinates and gene locations. c. Hi-M pairwise distance (PWD) matrix for nc14 embryos is shown at the top. Red and blue represent low and high spatial distance, respectively. Proximity frequency matrix with a cutoff of 250 nm from nc14 embryos is shown at the bottom. Pink and green represent high and low proximity frequency, respectively. d. Cumulative proximity frequency versus cutoff distance curve for class I IBPs barcodes (green) and control regions (yellow) for nc14 embryos. The gray shadow represents the delimitation of the threshold used to define co-localization (250 nm). A magnification of this zone is displayed in the inset. e. Violin plot distributions representing the frequency with which each insulator barcode interacts with any other insulator barcode in our oligopaint library for nc14 embryos (see Methods). Red line represents the mean. f. Normalized proximity frequency versus number of interacting partners for class I IBPs for nc14 embryos. g. Histograms of class I IBPs preferential interaction as a function of the number of interacting partners normalized by the number of pairwise interaction for nc14 embryos. Insulator barcodes are indicated on the left (I1 to I10). The color-scale represents the normalized frequency in log2-scale.

The proximity and PWD distance maps revealed multiple regions displaying preferential 3D spatial proximity (**Fig 3c**). Most regions in spatial proximity coincide with regions bound either on one or both anchors by insulators (yellow and green arrows respectively, **Fig. 3c**). Importantly, these preferential proximities were detected even at short spatial distances (e.g. below 100 nm) (**Fig. S3f**), and occurred between and across TAD borders, as well as within TADs. These results are therefore consistent with our previous findings that insulators are bound to regions displaying long-range contacts within and across TADs.

To quantify the frequency at which insulator-bound regions spatially co-localized in a population of single cells, we calculated the cumulative average proximity frequencies between insulator-bound regions and control regions (green and yellow barcodes, respectively) for different cutoff distances (**Fig. 3d**). At the cutoff distance used to calculate proximity maps (250 nm), insulator barcodes co-localized only in a small fraction of cells (~12%, **Fig. 3d** and inset). As expected, the proximity frequency monotonously increased with cutoff distance, but remained low for cutoff distance thresholds used in this and other studies (<250nm) ^25,30,38^. Thus, we conclude that colocalization between insulator-bound regions within and between TADs is rather infrequent, consistent with colocalization of insulator barcodes occurring only in a small proportion of cells (i.e. large cell-to-cell heterogeneity) or/and with colocalization being highly dynamic.

Next, we investigated the specificity of insulator barcode co-localizations by calculating the proximity frequency versus cutoff distance curve for non-insulator barcodes located at similar genomic distances (**Fig. 3d**, yellow curve). At a cutoff distance of 250 nm, non-insulator barcodes co-localized with a frequency only slightly higher than insulator barcodes (9% and 12%, respectively). Therefore, these results are consistent with insulators being involved in stabilizing long-range interactions that are already created by other mechanisms (e.g. polymer fluctuations). These infrequent interactions may be either short-lived or occur in only a subset of cells.

### Insulator barcodes most frequently co-localize in pairs

The existence of multiple focal peaks in the Hi-M matrix can be explained by two different models. On one hand, a low fraction of single cells can form rosette-like structures where multiple insulator-bound regions come together in space at once, as suggested by previous models ^39^. On the other hand, different combinations of insulator barcodes may co-localize at low-frequencies in a pairwise manner in single cells. In this case, the multiplicity of peaks in the Hi-M matrix would arise from ensemble averaging. To discern between these two models, we calculated how often insulator barcodes were proximal (i.e. at a distance ≤ 250 nm) to any other insulator barcode in single cells. This frequency was comparable for all the insulator barcodes investigated, and on average lower than 12% (**Fig. 3e**). Thus, in single cells, insulator barcodes interact with any other (genomically close) insulator barcode at low frequency.

Finally, to explore if these rare spatial encounters involved multiple insulator-bound regions, we calculated the proportion of clusters containing two (i.e. pairwise cluster) or multiple insulator barcodes (multi-way cluster). Clusters containing only two insulator targets were the most common in all cases (>65%) (**Fig. 3f**). Next, we calculated the frequency of multi-way clusters as a function of the number of barcodes in a cluster for all barcodes combined (**Fig. 3g**) or for each barcode independently at a distance ≤ 250 nm (**Fig. 3f**) and for different distance thresholds (Fig. S3h). The frequency of multi-way interactions rapidly decreased with the number of co-localizing targets. All in all, these results indicate that insulator-bound regions rarely form clusters with more than two insulators, and when they do, they contain only a very limited number of insulator-bound regions.

### Preferential interactions between class I insulators arise before TADs and transcription

Previous studies showed that preferential spatial proximity between certain *cis*-regulatory elements (e.g. enhancers and promoters) can be detected before the midblastula transition (MBT) occurring at nc14 ^30^, the nuclear cycle at which most zygotic genes get activated and when TADs first emerge ^20^. To investigate whether preferential interactions between insulator barcodes arose before these events, we compared Hi-M maps from nc12 and nc14 embryos by computing differential pairwise distance matrices (**Fig. 4a**). Notably, the overall structure of the *dpp* locus considerably changed during these two nuclear cycles as expected due to the emergence of TADs at nc14 (**Figs. 4a, S4a-c**). To better dissect how the proximity between insulator barcodes changed between nc12 and nc14, we calculated the proximity frequency versus cutoff distance curves for insulator and control regions (**Fig. 4b**). Insulator barcodes co-localized at similar frequencies in nc12 and nc14, but with a slightly reduced specificity at nc12, suggesting that preferred co-localization between insulator barcodes at the *dpp* locus is present as early as nc12.

**Figure 4.**
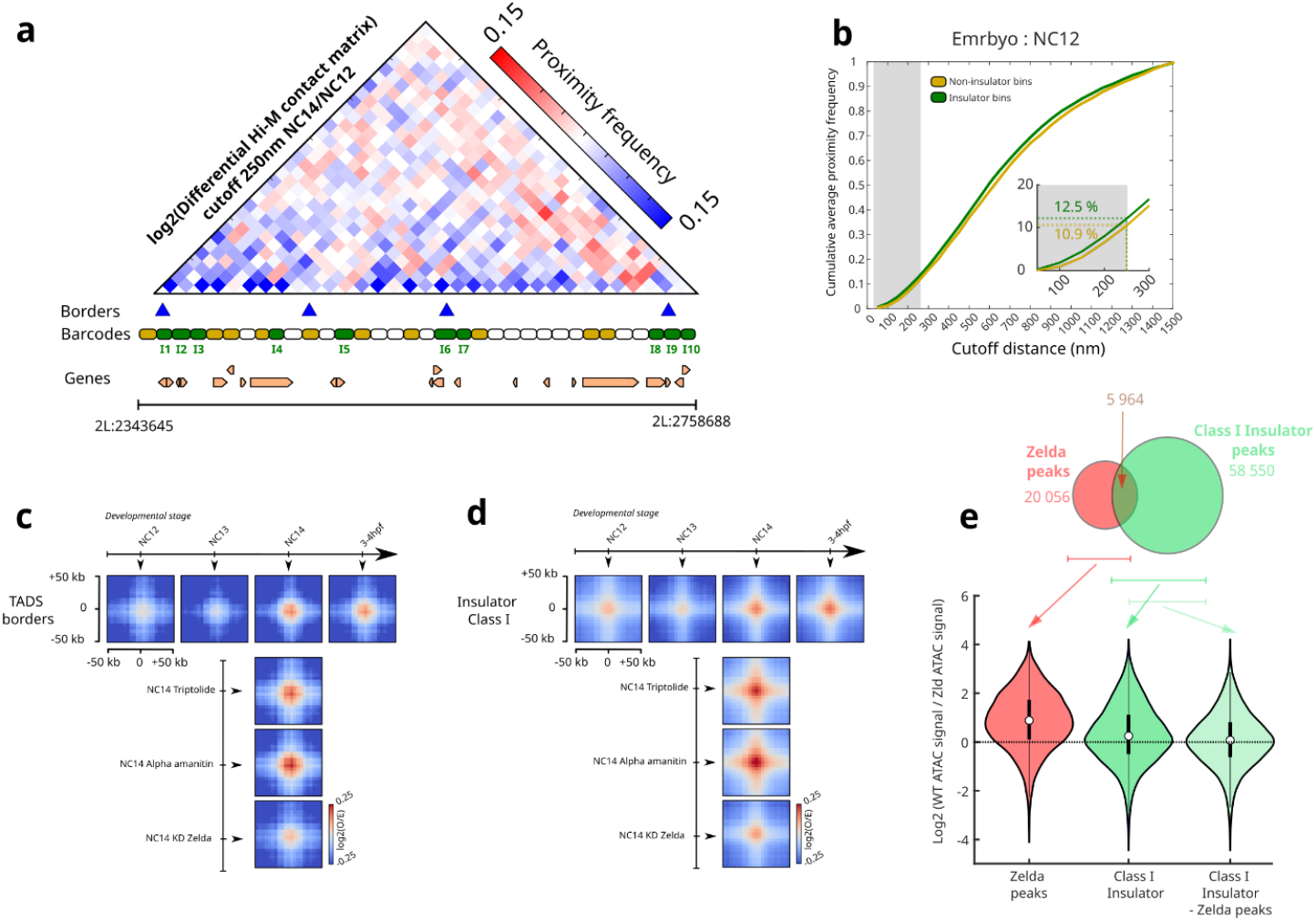
Border-Border interactions are formed at nc14, but non-border loops are gradually formed during development. a. Log2 differential pairwise Hi-M contact matrix between nc14 and nc12. Red and blue represent higher and lower contacts in nc14 as compared to nc12, respectively. b. Cumulative proximity frequency versus cutoff distance curve for class I IBPs barcodes (green) and control regions (yellow) for nc12 embryos. c. Aggregation Hi-C plots for TADs borders at different developmental stages (nc12, nc13, nc14, 3-4hpf) and for different biological conditions and treatment (nc14 triptolide-treated, nc14 alpha-amanitin-treated and nc14 knockdown of Zelda). d. Aggregation Hi-C plots for non-border class I IBPs at different developmental stages (nc12, nc13, nc14, 3-4hpf) and for different biological conditions and treatment (nc14 triptolide-treated, nc14 alpha-amanitin-treated and nc14 knockdown of Zelda). e. Pie plot representing the distribution of class I IBPs peaks and Zelda peaks (top panel). Violin plots showing the log2 ratio between the ATAC signal of wt embryo versus Zld-depleted embryos at Zelda regions (red) overlapping Zelda and class I IBPs peaks (dark green) and non-overlapping class I IBPs peaks (light green) (bottom panel).

To investigate whether this result could be generalized to other genomic regions, we performed APA analysis for nc12/nc13 (pre-ZGA), nc14 (ZGA) and 3-4 hours post fertilization (hpf) (post-ZGA). As expected, preferential interactions between TAD borders first appear at nc14 and are sustained thereafter (**Fig. 4c**), consistent with previous analyses ^20^. Thus, interactions between insulator-bound regions occupying TAD borders arise at the same time as TADs.

Next, we quantified the timing at which preferential interactions between non-border, insulator-bound regions emerged. For this, we performed APA analysis for non-borders for different developmental timings. Notably, we found that preferential interactions between non-border regions bound by insulators were already present in nc12 embryos for most class I IBPs (**Figs. 4d, S4e**), consistent with our Hi-M results at the *dpp* locus.

To further investigate the origin of these interactions, we performed APA analysis from nc14 embryos treated with triptolide and alpha-amanitin, two small-molecule inhibitors of RNA Pol II activity ^20^. Notably, preferential interactions between non-borders increased under these chemical perturbations (**Figs. 4d, S4f**). In contrast, our analysis shows that interactions between TAD borders were relatively undisturbed (**Fig. 4c**). The increase in interactions between non-border insulator-bound regions is consistent with enhanced inter-TAD interactions ^20^. As RNA Pol II activity in these embryos is inhibited before they are transcriptionally active, our result indicates that preferential interactions between non-border, insulator-bound regions seem to be independent of transcription.

Finally, to shed further light into the mechanism of interaction of non-border, insulator-bound regions, we performed APA analysis on embryos depleted in Zelda, a pioneering factor involved in establishing early accessibility of cis-regulatory elements ^40^. Surprisingly, preferential interactions between non-border, insulator-bound regions were overall unaffected in Zelda-depleted embryos (**Figs. 4d, S4e**), suggesting that binding of class I insulators to non-border regions does not require chromatin opening by Zelda. To test this hypothesis, we first calculated the fraction of class I IBP binding sites overlapping with Zelda sites. This analysis revealed that only ~10% of the class I IBP sites corresponded to Zelda sites (**Fig. 4e**). Next, we calculated chromatin accessibility at Zelda and class I IBP sites in wild-type and Zelda-depleted nc14 embryos. As expected, chromatin accessibility at Zelda-bound regions was severely reduced in Zelda-depleted embryos (**Fig. 4e**, red violin plot). In contrast, chromatin accessibility of class I insulators was not affected by Zelda-depletion (**Fig. 4e**, light green violin plot). Overall, these results are consistent with contacts between class I insulators not being affected by Zelda depletion (**Fig. 4d**), and indicate that this class of insulators rely on other means to access chromatin during early embryogenesis.

## Discussion

In this study, we applied bioinformatic analysis to investigate the role of *Drosophila* insulator binding proteins in the folding of the zygotic genome during early embryogenesis, and combined it with novel imaging-based chromosome conformation capture approaches to quantify the absolute frequency and specificity of pairwise and multiway chromatin interactions involving insulators.

Drosophila insulator proteins are highly enriched at TADs borders ^4,6,19,21^ and contribute to the insulation of TADs ^41–43^. Our bioinformatics analysis reveals that most preferential chromatin interactions genome-wide involve regions bound by class I insulators that do not involve TAD borders (>90%). This finding suggests that class I insulators are likely also involved in modulating interactions within TADs and across TAD boundaries. Members of the class I insulator family (e.g. BEAF-32) tend to co-localize with promoter regions ^44,45^ and tend to demarcate differentially-expressed genes ^46^, suggesting that class I insulators may play a role in modulating contacts between *cis*-regulatory modules within and between TADs. Direct promoter regulation and reduction in TAD insulation can only account for a minority (20%) of the genes downregulated upon depletion of BEAF-32 ^42^. Long-range, non-border chromatin interactions by class I IBPs appear before TADs and the onset of zygotic transition, suggesting that they contribute to defining pre-established topologies to demarcate *cis*-regulatory networks.

Zelda plays a central role in rendering the zygotic genome accessible ^47–50^. However, we found that interactions between class I IBPs at non-border regions are not affected by the depletion of Zelda. This surprising result may be explained by our finding that a significant portion of class I IBPs peaks (~90%) are open at early developmental cycles (e.g. nc12) but do not colocalize with Zelda, suggesting that other unidentified pioneering factors may be required to provide access to most class I IBPs.

Despite the genome-wide enrichment of IBPs at regions displaying 3D preferential interactions, the quantification of absolute proximity frequencies using Hi-M shows that insulator-bound regions (borders and non-borders) physically co-localize in space infrequently (~12%), and marginally more frequently than neighboring genomic regions (~9%). This observation is consistent with low proximity frequencies between TAD borders measured in S2 cells (~10%) ^35^. The low proximity frequencies between insulator-enriched regions are consistent with a recent study showing that depletion of insulators only partially weakens the strength of TAD borders ^42^, and with the overall absence of “focal loops’’ involving class I insulators in Hi-C contact maps ^4,6,20,29,51^. Genome-wide measurements of chromatin interactions will be required to extrapolate these results to long-range genomic distances.

The early discovery of insulator bodies led to the proposal that insulators mediate the formation of stable, rosette-like hubs involving multiple insulator-bound genomic regions ^52–55^. More recently, it was shown that CP190 and Su(HW) insulator bodies formed in cultured-cells under stress conditions exhibit liquid-liquid phase separation properties ^56^. This model predicts that genomically-close insulators should interact in space often, nucleating interactions between multiple partners. In contrast, we observed low-frequencies of pairwise proximities that rapidly decrease with the number of interacting partners (<5 % for 3-way interactions and <1 % for 4-way interaction). Therefore, these results do not provide support for a widespread role of stable insulator hubs or LLPS-mediated insulator bodies in the 3D organization of the *Drosophila* genome, at least in normal physiological conditions.

Instead, our observations support a model whereby insulators act by stabilizing short-lived 3D chromatin conformations between distant genomic regions ^57–59^ that mainly arise from the intrinsic dynamics of the chromatin polymer ^60^. The transient stabilization of 3D proximities promoted by the binding of insulators can provide a means to modulate the landscape of E-P contacts in 3D to either insulate promoters from the action of enhancers (e.g. enhancer blocking), or alternatively to promote transcription by bringing together actively transcribed regions. In this scenario, combinatorial binding of multiple insulator binding sites at single genomic locations ^19,42^ would provide a means to modulate the strength of the stabilization, to regulate it depending on cell type, and to enable a locus to time-share 3D interactions with multiple genomic locations in an asynchronous manner. Consistent with this concept, analyzing binding of RNAPII and polycomb members in mouse embryonic stem cell promoter-centered chromatin interactions using network measures such as bridgeness and betweenness centrality, it was suggested that RNAPII bound chromatin fragments would belong to multiple communities at once, whereas polycomb bound fragments appeared to participate in multiple interactions at once ^26^.

Direct measurements of residence times have, unfortunately, not been reported for class I *Drosophila* insulators. However, recent studies showed that GAF and mammalian CTCF can remain bound to their cognate chromatin sites for minutes ^61,62^, suggesting that insulator loops are dynamic ^58,63^. These data are consistent with a model whereby insulators help modulate the dynamics of specific interactions between distant cis-regulatory regions, but do not form stable scaffolds. These transient structures, however, are considerably more stable than the typical residence time of transcription factors (~10 seconds) ^64^. In this picture, insulators could promote transcription by stabilizing transient *cis*-regulatory interactions to allow for the rapid binding and unbinding of transcription factors, or rather repress transcription by promoting 3D conformations that prevent functional interactions.

Finally, the methods used in this manuscript to show that *Drosophila* insulators only moderately increase the frequency of border and non-border chromatin interactions may be used to investigate insulator mechanisms in other organisms.

## Methods

### Drosophila stocks and embryo collection

The *yw* fly stocks were maintained either in a 21°C room or in a 25°C incubator with a natural light-dark circadian cycle. Following a pre-laying period of 16-18 h in cages with yeasted apple juice agar plates, flies were allowed to lay eggs during 1.5 h on new plates. Layed embryos were then incubated at 25°C for an extra 2.5 h to reach the desired developmental stage. Embryos were collected and fixed as previously described ^37^.

### Hi-M libraries

Oligopaint libraries from a public database (http://genetics.med.harvard.edu/oligopaints), consisting of unique 35/45-mer sequences with genome homology. We selected 138 genomic regions of interest (barcodes) in the *dpp* locus (2L:2343645..2758688 BDGP *Release 6* + ISO1 *MT/dm6*). The coordinates of the targeted genomic regions are listed in Supplementary Data 2. The procedure to amplify a given library from the pool was previously described ^37^. For imaging, we used a combination of 4 barcodes to cover ~12kb, the list of positions of the barcodes are listed in Supplementary Table 2. The sequences of the adapters and labeled barcodes purchased from Integrated DNA Technology (IDT) are listed in Supplementary Data 1.

### Hybridization of Hi-M primary library

The ssDNA library is hybridized to the DNA as previously described ^37^. Briefly, embryos were rehydrated and permeabilized by sequential dilution of methanol with 0.1% Tween-20 PBS (PBT) : 90%MeOH; 70%MeOH; 50%MeOH; 30%MeOH; 100%PBT (5min each). Embryos were RNase A treated during 2h, permeabilized 1h with 0.5% Triton in PBS and rinsed with increased concentration of Triton/pHM buffer. pHM (pHM = 2X SSC, NaH_2_PO_4_ 0.1M pH = 7, 0.1% Tween-20, 50% formamide (v/v)) : 20%pHM; 50%pHM; 80%pHM ; 100%pHM (20 min each). Then, 225 pmols of ssDNA were diluted in 25μL of Fish Hybridization Buffer (FHB = 50% Formamide, 10% dextran sulfate, 2X SSC, Salmon Sperm DNA 0.5 mg/mL). The ssDNA and embryos were preheated at 80°C during 15 minutes in separated tubes. The supernatant of the embryo’s tube (pHM) is removed and the 25μL of FHB containing the ssDNA is added. Next the mixture is transferred in a PCR-tube and deposited in the thermomixer set at 80°C. Immediately, the thermomixer is set to decrease to 0.1°C/min until it reaches 37°C for an overnight incubation. The next day, the embryos were transferred to a new 1.5 mL eppendorf tube and washed two times at 37°C during 20 min with 50% formamide, 2X SSC. Next, embryos were sequentially washed at 37°C for 20 min with serial dilutions of formamide/PBT: 50% formamide / 2xSSC; 40% formamide / 2xSSC; 30% formamide / 70% PBT; 20% formamide / 80% PBT; 20% formamide / 80% PBT; 10% formamide / 90% PBT; 100% PBT. An additional crosslink step with PFA 4% was performed and labelled embryos were washed, resuspended in PBS and stored at −20°C for months until further use.

### Imaging system

Experiments were performed on a home-made imaging setup built on a RAMM modular microscope system (Applied Scientific Instrumentation) coupled to an improved microfluidic device, as the one described previously ^37^. Software-controlled microscope components, including camera, stages, lasers, needles, pump and valves, were run using Qudi-HiM, an homemade software developed in python ^65^ (RRID, record ID: SCR_022114). Embryos were imaged using an ×60 Plan-Achromat water-immersion objective (numerical aperture=1.2; Nikon) mounted on a closed-loop piezoelectric stage (Nano-F100, Mad City Labs Inc.). The Illumination was provided by three lasers (OBIS-405nm nm and Sapphire-LP-561nm from Coherent and VFL-0-1000-642-OEM1 from MPB communications Inc.) and the images were acquired using an sCMOS camera (ORCA Flash 4.0V3, Hamamatsu, Japan). A homemade autofocus system was used to correct for axial drift in real time using a 785nm laser (OBIS-785 datasets.

### Acquisition of Hi-M datasets

Embryos were aligned on a 2% agar:PBS pad, attached to a 1:10 poly(L-lysine):water coated coverslip and mounted into a FCS2® flow chamber (Bioptechs, USA). ~20-30 embryos were selected and imaged using two regions of interest (ROI 200×200μm^2^). Then, a mixture containing the fiducial adapter (25nM Atto-550 imager probe, 25nM of adapter to the reverse primer, 2× SSC, 40% v:v formamide) was injected in the chamber and let incubate for 15 min to allow complete hybridization on the primary FISH library. Embryos were washed for 10 min with a washing buffer solution (2× SSC, 40% v:v formamide) and for 5 min with 2×SSC before injecting 0.5 μg.ml-1 of DAPI in PBS to stain nuclei. Prior to imaging, the imaging buffer (1x PBS, 5% w:v glucose, 0.5mg/ml of glucose oxidase and 0.05mg/ml of catalase) was injected to reduce photobleaching of the fiducial barcode. A stack of images was acquired for DAPI and the fiducial tagged with Atto550 (z-step size of 250 nm and a total range of 20 μm) using 405 nm and 561 nm sequential illumination. Next, the sample was sequentially hybridized as follows. A solution containing the barcode and the imager oligo was injected (25nM Alexa-SS-647 probe, 25nM barcode, 2× SSC, 40% v:v formamide) and incubated for 15 min. Then, the embryos were washed with 1.5mL of washing buffer and with 1.5 mL of 2x SSC before injecting the imaging buffer. In each cycle, fiducials and readout probes were sequentially imaged with 561 nm and 647 nm excitation lasers. Finally, samples were washed with 1 mL of 2× SSC for 5 min before a new hybridization cycle started. Further details can be found on our previously published protocol ^37^.

### Image processing

DCIMG files were converted to TIFF using proprietary software from Hamamatsu. TIFF images were then deconvolved using Huygens Professional 21.04 (Scientific Volume Imaging, https://svi.nl). The analysis was performed using our pyHiM analysis pipeline (https://pyhim.readthedocs.io/en/latest/). Briefly, images were first z-projected using either sum (DAPI channel) or maximum intensity projections (barcodes, fiducials). Fiducial images from each hybridization cycle were used to register barcode images using global and local registration methods. Next, barcode images were segmented in 3D using stardist ^66^ and the positions of the centers of barcodes were detected with subpixel resolution using Big-FISH (https://github.com/fish-quant/big-fish) ^67^. Finally, pairwise distance matrices were calculated for each single nucleus. From the list of pairwise distance maps, we calculated the proximity frequencies as the number of nuclei in which pairwise distances were within 250 nm normalized by the number of nuclei containing both barcodes.

### Multi-way proximity frequency analysis

The proportion of multi-way contacts is calculated from single nucleus proximity frequency matrices as described previously ^68^. Briefly, we counted the number of multi-way contacts where the selected anchor barcode was interacting with other partners within a 250 nm radius. These values were normalized by the number of pairwise interactions for each anchor.

### Chip-Seq data processing

Insulator proteins ChIP-Seq fastq files were downloaded from Gene Expression Omnibus (GEO). The quality of the reads was estimated with FastQC ^69^ (0.11.7). Sequencing reads were aligned to the reference drosophila melanogaster genome assembly (dm6) using Burrows-Wheeler Aligner ^70^ (0.7.17-r1188) with default parameters. Finally, peak calling was performed using MACS2 ^71^ (2.2.7.1) with default parameters. The accession number of the data used in this study are listed in Supplementary Table 1.

### Chromatin Assortativity

In order to build networks needed for Chromatin Assortativity, Hi-C ^20^ contact matrices were used with a 5-kb resolution. From such matrices, significant interactions were called using Chromosight ^28^ (1.3.3) with default parameters. For each stage or condition, a specific genome scale chromatin network was built where nodes are genomic fragments and edges are significant interactions between two fragments. Then, chromatin networks were loaded on R and ChIP-seq peaks were used as features assigned to nodes using the ChAseR R package ^72^ (0.0.0.9) to calculate chromatin assortativity. For each feature, 1000 randomized networks preserving genomic distances and corresponding chromatin assortativity values were computed.

The chromatin assortativity Z-Scores calculation is given by the following formula :

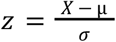

where *X* is the feature chromatin assortativity value, *μ* is the randomizations ChAs average and *σ* is the standard deviation from the randomization distribution.

The calculation of the Z-score allows estimating the significance of the assortativity values with respect to the assortativity expected based purely on correlation of feature values along the linear genome. For example, domains of a feature that span multiple bins in the Hi-C matrix are more likely to produce high ChAs values, but this does not imply the importance of the 3D contacts.

Cytoscape (3.8.0) was used for chromatin networks visualization.

### Aggregate peak analysis

ChIP-seq peaks were used to extract a list of regions bound by a set of putative factors. Next, for each autosomal arm, we computed the log2(Observed/Expected) Hi-C contact value by calculating the average contact frequency for all the combinations of regions separated by a certain genomic distance on a 5kb-Hi-C dataset ^20^. We then selected the interaction between the regions bound by the set of factors on the distance normalized Hi-C dataset. The distribution of log2(O/E) between all pairwise combinations of regions bound by the investigated factors was then displayed in a violin plot or divided into equal-size groups depending either on protein occupancy or insulation score and displayed in Hi-C aggregate plots.

### Hi-C aggregate plot analysis

Hi-C aggregate plots were performed using a homemade analysis pipeline developed in MATLAB Release R2019b (The MathWorks, Inc., Natick, United States). The distance-normalized sub-matrices over a window of 50kb surrounding the peak anchor were extracted. Finally, the aggregate plots were then created by averaging all of the sub-matrices together.

## Acknowledgements

This project was funded by the European Union’s Horizon 2020 Research and Innovation Program (Grant ID 724429) (M.N.). We acknowledge the Bettencourt-Schueller Foundation for their prize ‘Coup d’élan pour la recherche Française’, and the Drosophila facility (BioCampus Montpellier, CNRS, INSERM, Univ Montpellier, Montpellier, France). The CBS is a member of the France-BioImaging, a national infrastructure supported by the French National Research Agency (ANR-10-INBS-04-01). O.M. was supported by an FRM PhD fellowship. F. R. and V.P. were supported by Fondation Toulouse Cancer Santé and Pierre Fabre Research Institute as part of the Chair of Bioinformatics in Oncology of the CRCT.

## Author Contributions

O.M., V.P. and M.N. conceived the study and the design. O.M. acquired the data. O.M., F.R., and J.G. analyzed the data. O.M., J.-B.F., and F.R. wrote the software. J-B.F. built the microscope. O.M., F.R., V.P. and M.N. interpreted the data. M.N., O.M., and V.P. wrote the manuscript. M.N. and V.P. supervised the study and acquired funds.

