## Supplementary Data for "3D chromatin interactions involving *Drosophila* insulators are infrequent but preferential and arise before TADs and transcription"

### Table of contents

**Table of contents**	**2**

**Supplementary Figures**	**3**

Supplementary Figure S1.	3

Supplementary Figure S2.	4

Supplementary Figure S3.	5

Supplementary Figure S4.	6

**Supplementary Tables and Data**	**8**

### Supplementary Figures

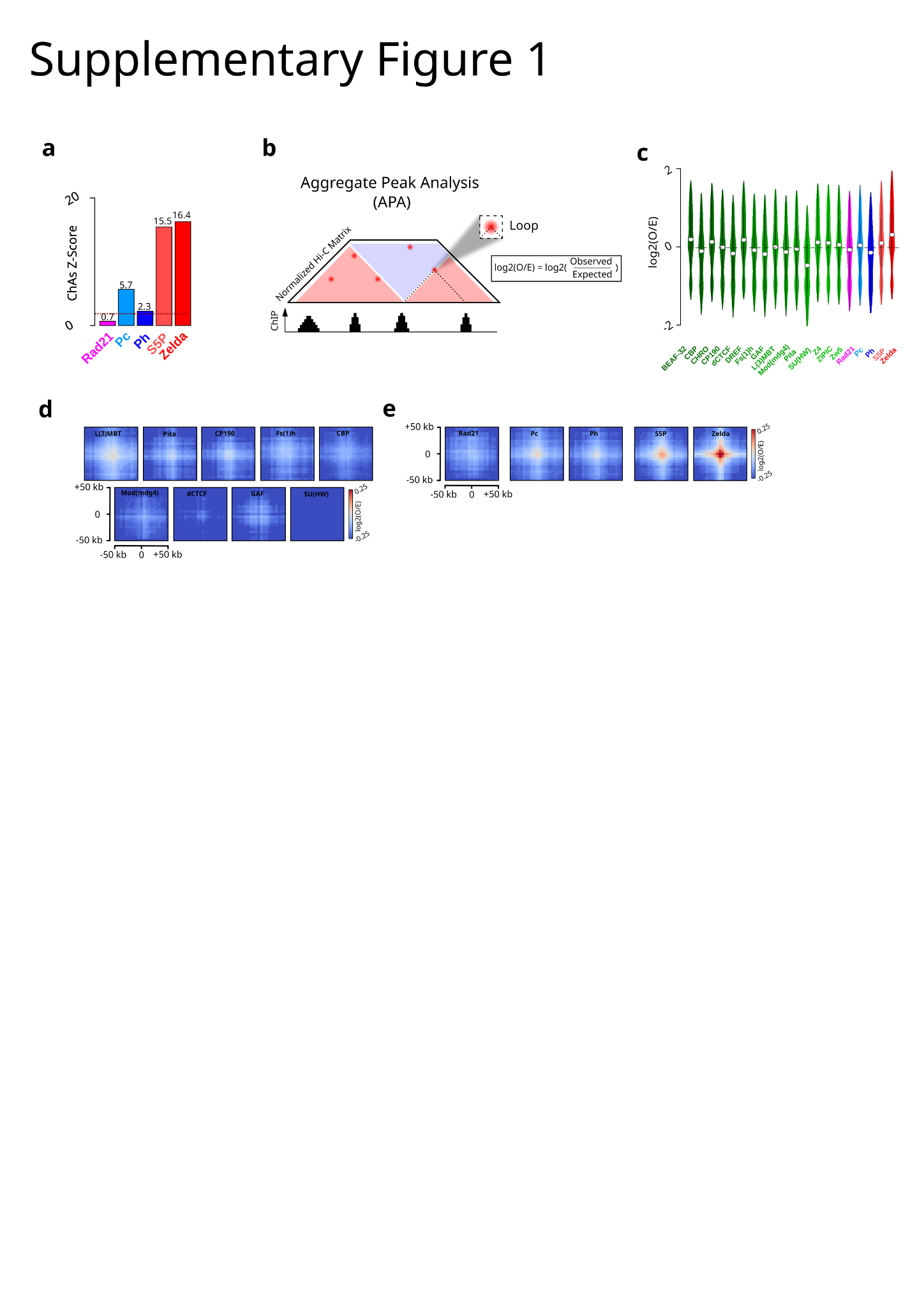

#### **Supplementary Figure S1.**

a. Barplot illustrating the ChAs-Z-Scores for the cohesin subunit (Rad21), Polycomb group proteins (Pc, Ph), RNA polymerase II (RNAPII CTD phospho-Ser5) and the pioneering factors (Zelda) in the nc14 chromatin network classified by alphabetical order.

b. Cartoon illustrating Aggregate Peak Analysis (APA) of chromatin binding factors in distance normalized Hi-C matrix.

c. Violin plots illustrating the O/E-Score for 15 IBPs, the cohesin subunit (Rad21), Polycomb group proteins (Pc, Ph), RNA polymerase II (RNAPII CTD phospho-Ser5) and the pioneering factors (Zelda) in the nc14 chromatin network classified by alphabetical order.

d. Aggregation Hi-C plots for nc14 embryos for class II IBPs that do not display a positive log2(O/E) : L(3MBT), Pita, CP190, Fs(1)h, CBP, Mod(mdg4), dCTCF, GAF, SU(HW).

e. Aggregation Hi-C plots for nc14 embryos for the cohesin subunit (Rad21), Polycomb group proteins (Pc, Ph), RNA polymerase II (RNAPII CTD phospho-Ser5) and the pioneering factors (Zelda).

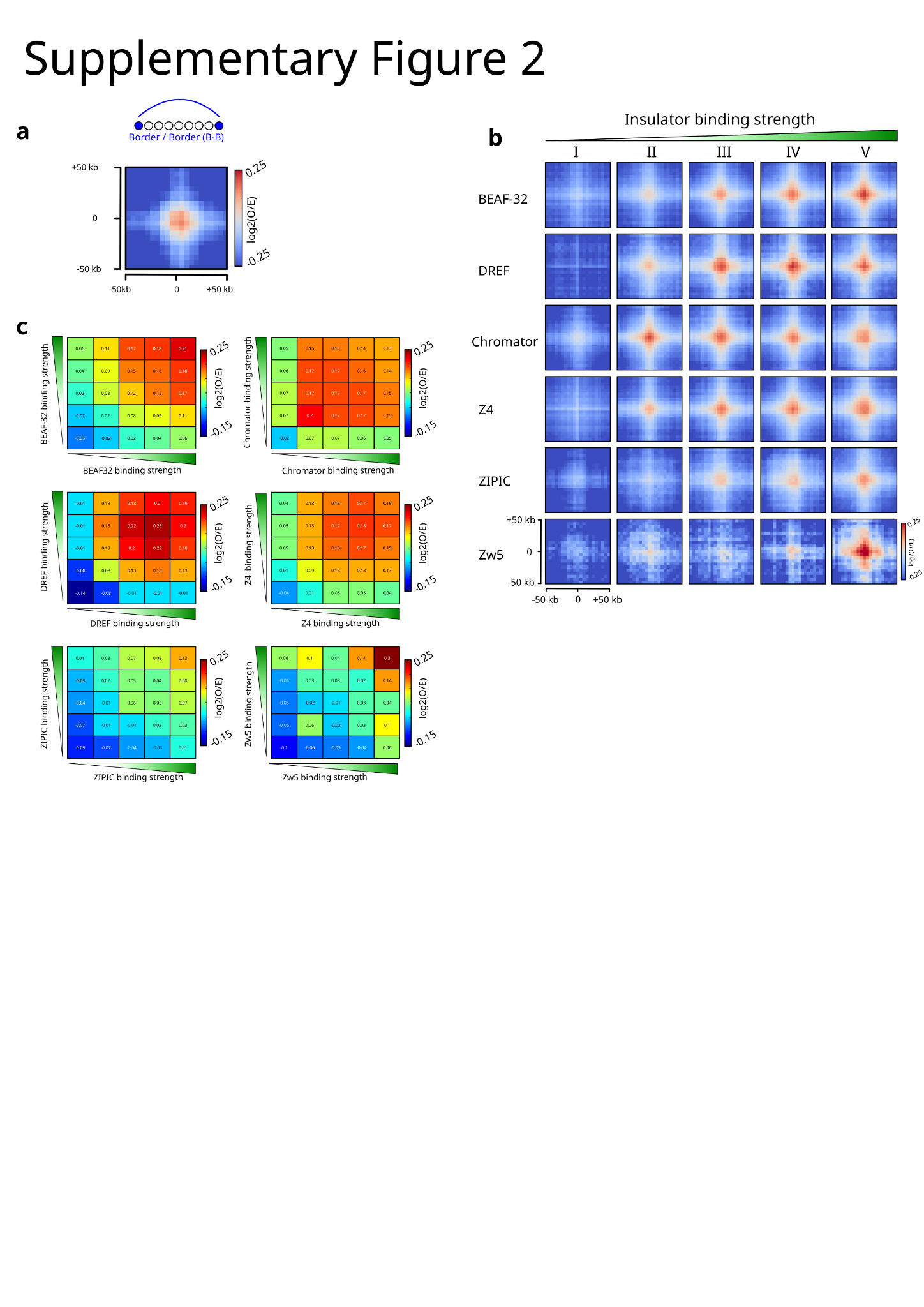

#### **Supplementary Figure S2.**

a. Aggregation Hi-C plots based Hi-C data for identified TADs borders in nc14.

b. Aggregation Hi-C plots for each individual protein of the class I IBP group in nc14 embryos classified by increasing level of binding.

c. log2(O/E) for each individual protein of the class I IBP group in nc14 ranked by increasing insulator enrichment in nc14 embryos.

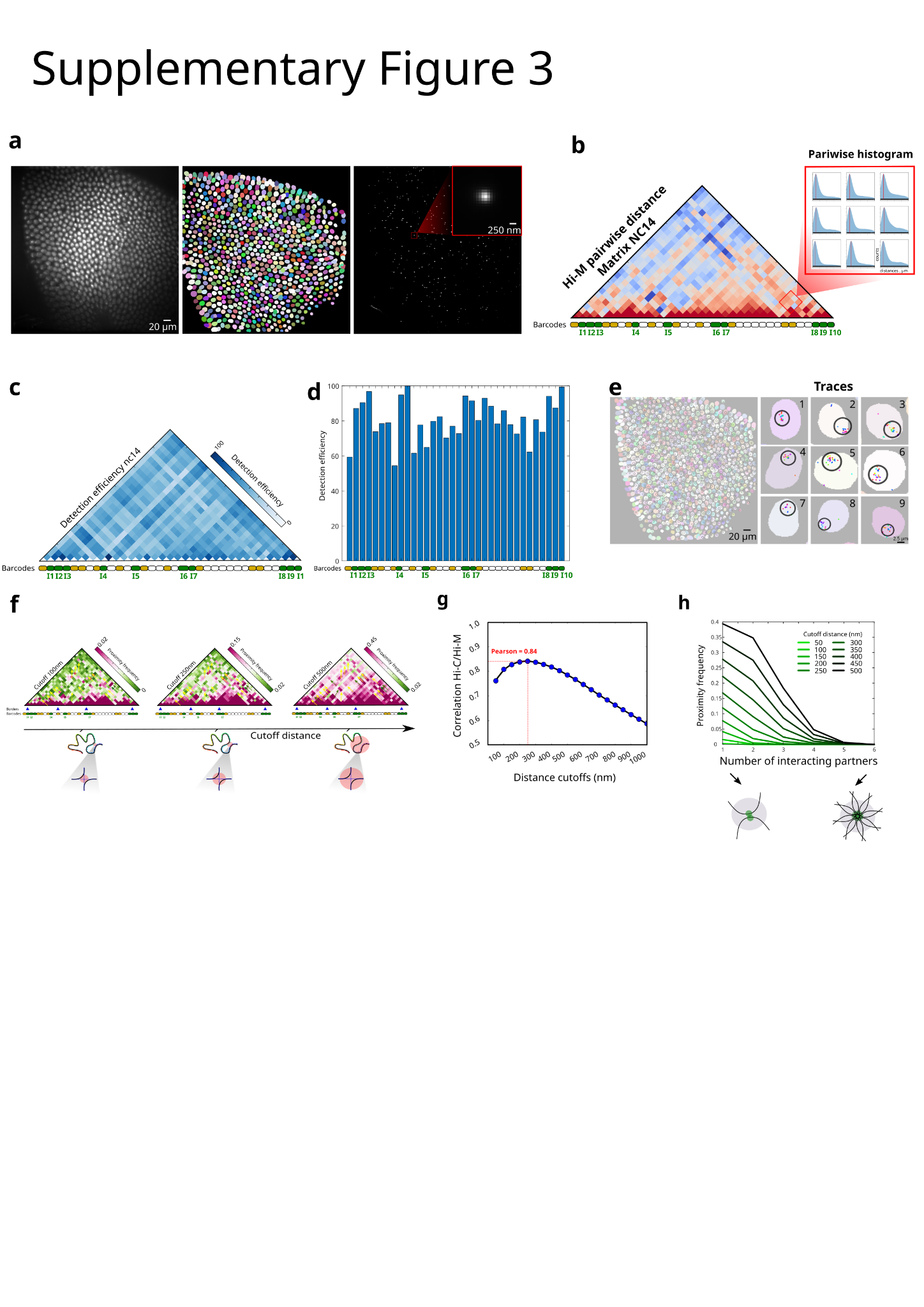

#### **Supplementary Figure S3.**

a. Left: grayscale image of DAPI-stained nuclei from nc14 Drosophila embryo. Middle : segmented DAPI masks after nuclei segmentation. Right : maximum intensity projection of the fluorescence signal from a single barcode in the same field of view.

b. Hi-M pairwise distance (PWD) matrix for nc14 embryos. The zoom shows the histogram of the pairwise distribution of distances.

c. Detection efficiency matrices normalized by the most detected barcode pair. Dark blue and white represent high and low detection efficiency respectively.

d. Barplot representing the Detection efficiency for all barcodes normalized by the most detected barcode.

e. Example of segmented barcodes in traces in the same field. The Color code indicates different barcodes found in a trace.

f. Hi-M proximity frequency matrices generated with various cutoff distances for nc14 embryos. From left to right the cutoffs considered are 100, 200, 500 nm.

g. Pearson correlation coefficient between interpolated Hi-C contact map and Hi-M proximity frequency maps generated with different cutoff distances. The higher correlation is reached with a cutoff of 250 nm.

h. Normalized proximity frequency versus number of interacting partners for class I IBPs with various cutoff distances for nc14 embryos.

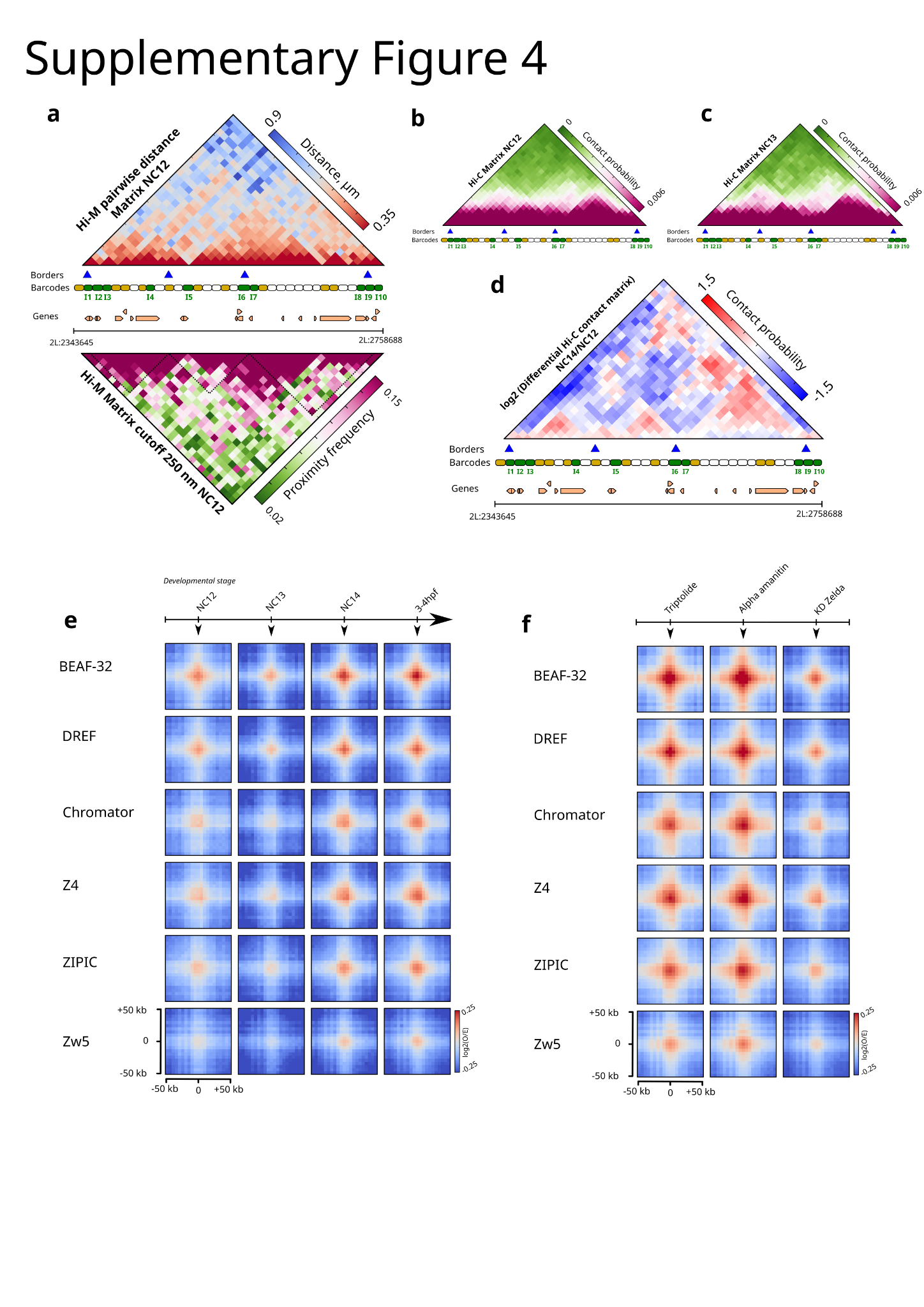

#### **Supplementary Figure S4.**

a. Top : nc12 pairwise distance matrix along the dpp locus (2L:2343645-2758688) in Drosophila melanogaster (dm6). Bottom : proximity frequency matrix with a cutoff of 250 nm.

b. Nc12 Hi-C matrix along the *dpp* locus (2L:2343645-2758688) in Drosophila melanogaster (dm6). Purple and green represent high and low contact probability respectively.

c. Nc13 Hi-C matrix along the *dpp* locus (2L:2343645-2758688) in Drosophila melanogaster (dm6). Purple and green represent high and low contact probability respectively.

d. Log2 differential pairwise Hi-C contact matrix between nc14 and nc12. Red and blue represent higher and lower contacts in nc14 as compared to nc12, respectively.

e. Aggregation Hi-C plots for each individual protein of the class I IBPs group at different developmental stages (nc12, nc13, nc14, 3-4hpf).

f. Aggregation Hi-C plots for each individual protein of the class I IBPs for different biological conditions (nc14 triptolide-treated, nc14 alpha-amanitin-treated and nc14 knockdown of Zelda).

### Supplementary Tables and Data

**Supplementary_Tables.xlsx**

SUPPLEMENTARY TABLE 1. List of publicly available data used in this study.

SUPPLEMENTARY TABLE 2. List of genomic positions of the barcodes used in this study.

SUPPLEMENTARY TABLE 3. List of primers for library amplification used in this study.

**Supplementary_Data_1.xlsx** contains the list of sequences of imaging (io), adapter oligos and barcodes used in this study.

**Supplementary_Data_2.xlsx** contains the list of sequences of primary Hi-M probes.
